# Upstream alternative polyadenylation in *SCN5A* produces a short transcript isoform encoding a mitochondria-localized NaV1.5 N-terminal fragment that influences cardiomyocyte respiration

**DOI:** 10.1101/2024.08.09.607406

**Authors:** Nathan H. Witmer, Jared M. McLendon, Colleen S. Stein, Jin-Young Yoon, Elena Berezhnaya, John W. Elrod, Barry L. London, Ryan L. Boudreau

**Affiliations:** Department of Internal Medicine, Fraternal Order of Eagles Diabetes Research Center, Abboud Cardiovascular Research Center, Carver College of Medicine, University of Iowa, Iowa City, IA; Molecular Medicine Graduate Program, Carver College of Medicine, University of Iowa, Iowa City, IA; Department of Pharmaceutical Sciences and Experimental Therapeutics, College of Pharmacy, University of Iowa, Iowa City, IA; Aging and Cardiovascular Discovery Center, Department of Cardiovascular Sciences, Lewis Katz School of Medicine, Temple University, Philadelphia, PA

## Abstract

*SCN5A* encodes the cardiac voltage-gated Na+ channel, NaV1.5, that initiates action potentials. *SCN5A* gene variants cause arrhythmias and increased heart failure risk. Mechanisms controlling NaV1.5 expression and activity are not fully understood. We recently found a well-conserved alternative polyadenylation (APA) signal downstream of the first *SCN5A* coding exon. This yields a *SCN5A-short* transcript isoform expressed in several species (e.g. human, pig, and cat), though rodents lack this upstream APA. Reanalysis of transcriptome-wide cardiac APA-seq and mRNA-seq data shows reductions in both upstream APA usage and short/full-length *SCN5A* mRNA ratios in failing hearts. Knock-in of the human *SCN5A* APA sequence into mice is sufficient to enable expression of *SCN5A*-short transcript, while significantly decreasing expression of full-length *SCN5A* mRNA. Notably, *SCN5A*-short transcript encodes a novel protein (NaV1.5-NT), composed of an N-terminus identical to NaV1.5 and a unique C-terminus derived from intronic sequence. AAV9 constructs were able to achieve stable NaV1.5-NT expression in mouse hearts, and western blot of human heart tissues showed bands co-migrating with NaV1.5-NT transgene-derived bands. NaV1.5-NT is predicted to contain a mitochondrial targeting sequence and localizes to mitochondria in cultured cardiomyocytes and in mouse hearts. NaV1.5-NT expression in cardiomyocytes led to elevations in basal oxygen consumption rate, ATP production, and mitochondrial ROS, while depleting NADH supply. Native PAGE analyses of mitochondria lysates revealed that NaV1.5-NT expression resulted in increased levels of disassembled complex V subunits and accumulation of complex I-containing supercomplexes. Overall, we discovered that APA-mediated regulation of *SCN5A* produces a short transcript encoding NaV1.5-NT. Our data support that NaV1.5-NT plays a multifaceted role in influencing mitochondrial physiology: 1) by increasing basal respiration likely through promoting complex V conformations that enhance proton leak, and 2) by increasing overall respiratory efficiency and NADH consumption by enhancing formation and/or stability of complex I-containing respiratory supercomplexes, though the specific molecular mechanisms underlying each of these remain unresolved.

## INTRODUCTION

Defects in heart function and arrhythmias are known to increase the risk of sudden cardiac death and often progress to heart failure (HF)^1^. Despite advances in treatment and prevention strategies, HF remains the world’s leading cause of death and affects ∼2% of people globally^2^. A host of underlying molecular mechanisms coincide with deterioration of cardiac function and arrhythmic death risk, including shifts in gene expression and ion channel dysregulation^3^. While the primary drivers (specific ion channels) mediating the cardiac action potential are known, there remains a need to thoroughly characterize the expansive regulatory mechanisms controlling these genes/proteins, as well as the many others that are central to HF onset and progression.

*SCN5A* encodes the pore-forming subunit of the voltage-gated Na+ channel, NaV1.5, that is responsible for inward Na+ current during the fast depolarization phase of the cardiac action potential^4^. Fine-tuned NaV1.5 activity is required for normal heart function^5^, and several genetic variants in *SCN5A* cause inherited arrhythmias (e.g. Brugada syndrome and long-QT syndrome) and dilated cardiomyopathy (DCM)^6,7^. Prior research has focused on defining NaV1.5 functional properties, effects of disease-causing variants, and therapeutic approaches to alter NaV1.5 activity^4,8–10^. Beyond that, some studies have also found decreased *SCN5A*/NaV1.5 expression and dysregulated function in non-arrhythmogenic HF, spurring investigations beyond conduction^11,12^. Along these lines, we previously reported that lower *SCN5A* expression associates with increased HF patient mortality, which was surprisingly linked to worsening ejection fraction as opposed to arrhythmic sudden cardiac death^13^. NaV1.5 channel function is regulated at several points during its lifecycle (e.g. intracellular trafficking, membrane anchoring, and degradation^14^, post-translational modification^15^, and interactions with metabolites^16–19^). Efforts to understand mechanisms controlling *SCN5A* expression have identified and begun to characterize several alternatively spliced transcripts^20,21^, transcriptional regulators^22–24^, RNA-binding proteins^25,26^, and microRNAs^13,25,27^. However, further characterization of *SCN5A*/NaV1.5 expression and function are needed to understand its role in cardiac biology and disease and beyond.

Alternative polyadenylation (APA) is an RNA regulatory mechanism that generates distinct 3’ transcript ends through differential usage of polyadenylation sites. APA involves ∼85 distinct regulatory proteins that generate APA signatures in ∼70% of human genes^28,29^. Notably, APA can occur in upstream introns to produce truncated protein isoforms^30^ and differential APA has been described in several conditions such as proliferation, differentiation, and membrane excitability^31–36^. Transcriptome-wide maps of APA events in human heart tissues support that APA is dynamically altered in HF. Global shortening of 3’-UTR lengths has also been observed in hypertrophied mouse hearts^37,38^. Despite these findings, the overall regulation and biological consequences of APA in the heart remain largely underexplored, both globally and at the single gene level.

Herein, we report our serendipitous discovery of a novel APA signal in the *SCN5A* locus that regulates the expression of full-length *SCN5A* mRNA and produces a short transcript isoform that encodes an N-terminal fragment of NaV1.5 (NaV1.5-NT). Interestingly, our initial characterizations support that NaV1.5-NT localizes to mitochondria and influences cardiomyocyte respiration.

## RESULTS

### An alternative polyadenylation signal downstream of *SCN5A* exon 2 generates a truncated transcript that is downregulated in failing human hearts

While exploring the *SCN5A* gene structure and expression patterns on the UCSC Genome Browser^39^, we noticed the presence of RNA-seq reads (GTEx human heart RNA-seq tracks) extending into the intronic region downstream of the first coding exon (exon 2), and these reads appeared to dissipate near a canonical polyadenylation signal sequence (AATAAA; **Fig. 1A,B**). We also observed this pattern in several myocyte-like cell lines (AC16, Rh30, and iPS-CMs) and in cardiac RNA-seq data from other species (e.g. cat, pig, and monkey; data not shown). Notably, this upstream polyA signal sequence is reasonably well-conserved across mammals (present in 27/62 species) but is not present in mice or rats (**Fig. 1B**). Targeted 3’-end polyA-seq^37^ done in nonfailing and failing human heart tissues shows a convincing transcript terminus mapping ∼30 bases downstream of the polyA signal in all samples analyzed, consistent with canonical 3’-end mRNA processing at this site^40^ (**Fig. 1A, Supplementary** Fig. 1). Taken together, these data indicate that this previously overlooked polyA signal generates a short *SCN5A* transcript that we refer to as *SCN5A-short*. To assess whether *SCN5A* APA might be altered in HF, we performed targeted re-analysis of the Creemers et al. human cardiac 3’-end RNA-seq data, which indicated a trending reduction in the ratio of *SCN5A-short*/full-length polyA usage in patients with DCM (versus nonfailing controls, p<0.09, n≥5/group; **Fig. 1C**). In addition, re-analysis of human cardiac RNA-seq data from two larger cohorts^41,42^ provides further support that *SCN5A-short* transcript expression (relative to full-length *SCN5A mRNA*) is decreased in failing hearts (DCM and ischemic cardiomyopathy [ICM], versus nonfailing controls, p<0.05, **Fig. 1D,E**).

**Figure 1.**
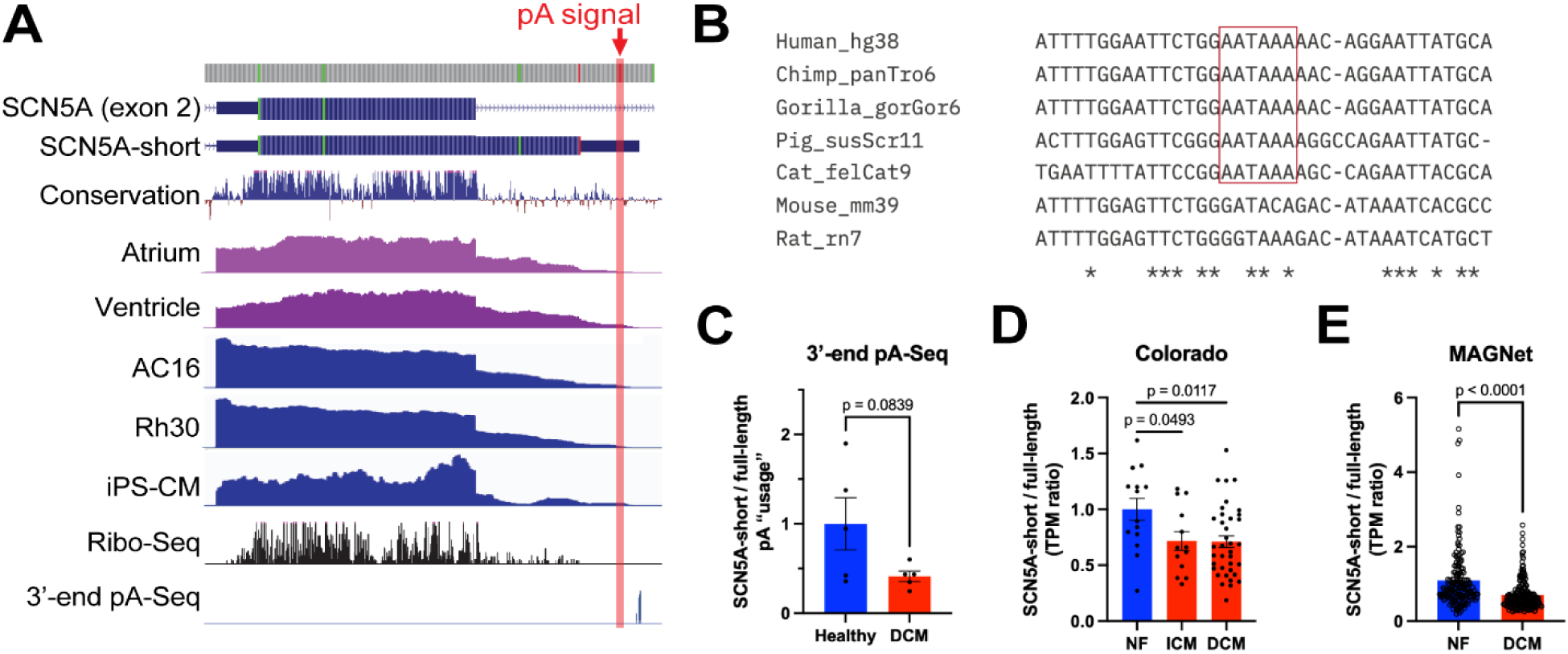
An alternative polyadenylation signal downstream of SCN5A exon 2 generates a truncated transcript that is downregulated in failing human hearts. A) Screenshots of the *SCN5A* exon 2 locus from the UCSC Genome Browser (hg38) with NCBI RefSeq gene annotations, GTEx RNA-seq (representative human atrial and left ventricular), and GWIPS-viz Ribo-seq tracks are shown. Representative RNA-seq from AC16, Rh30, and iPS-cardiomyocyte cell lines and 3’-end polyA-seq from human heart tissue are also shown. Red box indicates the polyadenylation signal sequence (AATAAA), which leads to the production of a truncated transcript isoform, *SCN5A-short*. **B)** Clustal-Omega alignment of the alternative polyadenylation signal in selected mammals, showing its conservation in several species but absence in mice or rats. **C)** Targeted analysis of 3’-end polyA-seq in healthy versus dilated cardiomyopathy (DCM) human heart samples. The relative ratio of reads at the upstream polyA site to the full-length polyA site is plotted as mean ± SEM, and p-value was determined by an unpaired t-test. **D)** Re-analysis of human heart RNA-seq data (Colorado study) comparing the relative expression of short to full-length *SCN5A* transcript expression in non-failing versus ischemic cardiomyopathy (ICM) and dilated cardiomyopathy (DCM). The relative TPM ratio is plotted as mean ± SEM, and p-value was determined by a one-way ANOVA with Dunnett’s post-hoc. **E)** Re-analysis of human heart RNA-seq data from the MAGNet (UPenn) study comparing the relative expression of short to full-length *SCN5A* transcript expression in non-failing versus dilated cardiomyopathy (DCM). The relative TPM ratio is plotted as mean ± SEM, and p-value was determined by an unpaired t-test.

### The *SCN5A* alternative polyadenylation signal influences full-length *SCN5A* mRNA expression

To assess whether the upstream polyA site influences full-length *SCN5A* mRNA and NaV1.5 protein expression, we generated a “humanized” knock-in mouse model (*SCN5A*-alt-pA KI). For this, we replaced the mouse intronic region with the human *SCN5A* sequence harboring the polyA signal and some additional surrounding sequence context (**Fig. 2A**). RT-qPCR done on wildtype (WT) and *SCN5A*-alt-pA KI mouse heart tissue samples indicates that the alternative polyA site is used in KI mice to generate *SCN5A-short* (**Fig. 2B**). Additional RT-qPCR analyses also revealed that full-length *SCN5A* mRNA expression is significantly lower in alt-pA-KI mouse hearts (versus WT controls, n≥4/group, p<0.05; **Fig. 2C**). However, full-length NaV1.5 protein expression is unchanged (**Fig. 2D**). Overall, these findings support that the *SCN5A* APA signal is utilized and can influence full-length *SCN5A* mRNA expression, while potential compensatory regulatory mechanisms may contribute to maintaining normal levels of NaV1.5.

**Figure 2.**
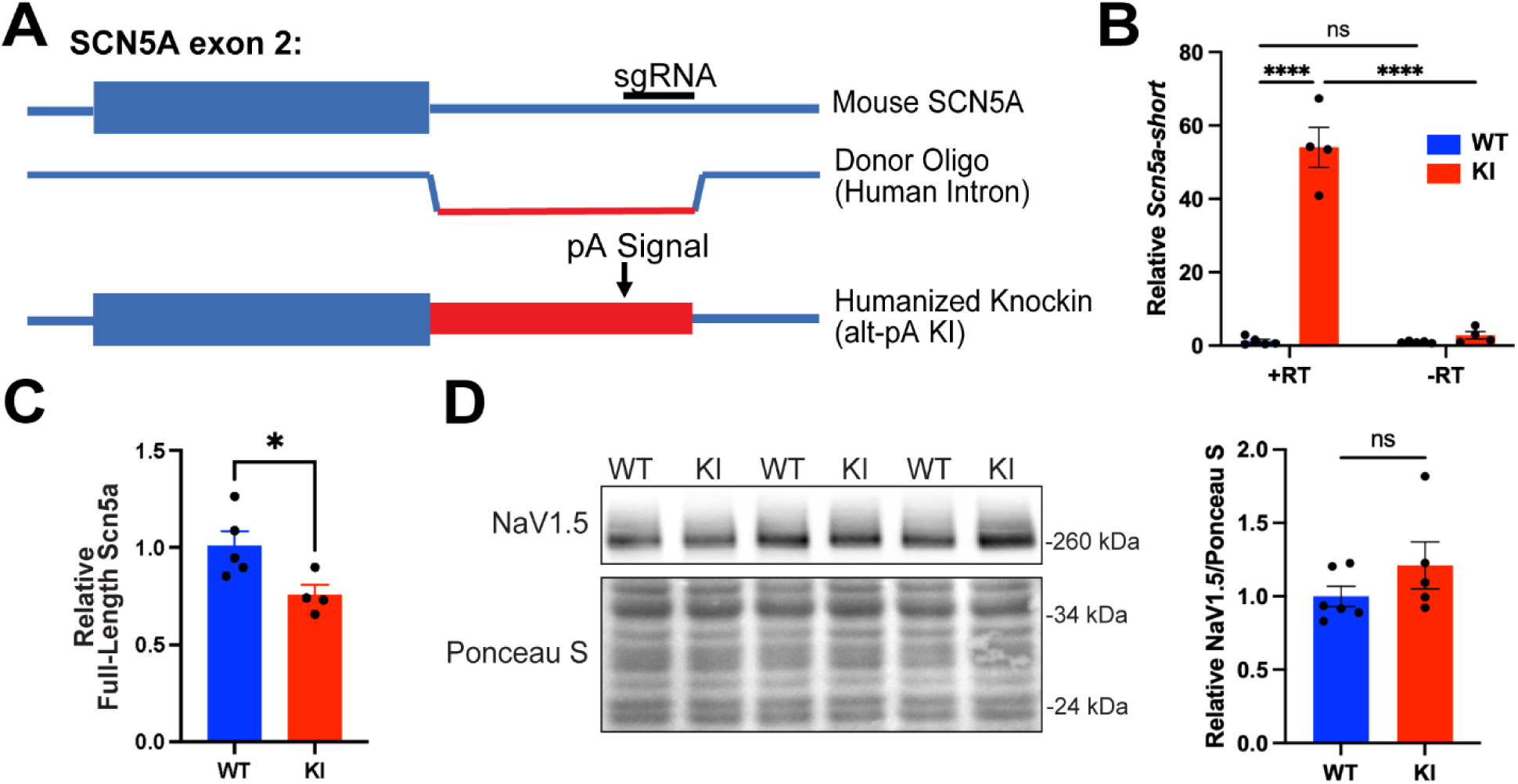
The *SCN5A* alternative polyadenylation signal regulates full-length *SCN5A* mRNA expression. A) Schematic of how the *SCN5A*-alt-pA knock-in mouse line was generated. CRISPR/Cas9 technology was employed to replace the mouse intronic region with the human version, which carries the alternative polyA signal sequence. Red region indicates the edited sequence. **B)** RT-qPCR for *SCN5A-short* transcripts in wildtype (WT) versus *SCN5A*-alt-pA knock-in (KI) mice, with or without reverse transcriptase (RT), supporting that this polyA signal can generate *SCN5A-short*. Relative expression is plotted as mean ± SEM, and p-value was determined by two-way ANOVA with Fisher’s LSD post-hoc (****p<0.0001). **C)** RT-qPCR for full-length *SCN5A* transcripts in WT versus KI mice, indicating alternative polyadenylation regulates its expression. Relative expression is plotted as mean ± SEM, and p-value was determined by an unpaired t-test (*p<0.05). **D)** Western blot and quantitation of NaV1.5 protein expression in WT and KI mouse heart lysates. Relative expression is plotted as mean ± SEM, and p-value was determined by an unpaired t-test.

### *SCN5A-short* transcript generates a novel protein corresponding to the N-terminus of NaV1.5

In addition to APA-mediated regulation of full-length *SCN5A* expression, the truncated transcript isoform could generate a novel protein composed of an N-terminus identical to NaV1.5 (residues 1-91) and a unique C-terminus derived from “intronic” sequence (residues 92-134), termed NaV1.5-NT (**Fig. 3A**). To begin characterizing the expression and biochemistry of this protein, we first assessed its potential expression in neonatal rat cardiomyocytes (NRCMs) using an adenovirus (Ad-NaV1.5-NT) and subjected cell lysates to western blotting using a commercial NaV1.5 N-terminal antibody (recognizes aa 42-70) and a custom antibody that we generated to recognize the unique C-terminal region (aa 109-123). This revealed the presence of three unique bands (migrating at ∼17-20 kDa) that derive from the NaV1.5-NT transgene (**Fig. 3B**). In similar fashion, we generated a cardiomyocyte-specific AAV (AAV9-NaV1.5-NT, driven by the human cardiac troponin promoter) to express NaV1.5-NT in mouse heart tissues. Three weeks after AAV administration in mice, cardiac tissue lysates were assessed by western blot. Blotting with the commercial NaV1.5-NT antibody revealed unique NaV1.5-NT-derived bands running at ∼17 kDa and 34 kDa (possible SDS-resistant dimers), the former of which appears to migrate at the same size as a background band in vehicle-injected controls. Notably, 17 kDa NaV1.5-NT is co-immunoreactive with our custom antibody, which does not appear to detect 34 kDa NaV1.5-NT (**Fig. 3C**); it is possible the antibody epitope is not accessible in the dimer. We also assessed NaV1.5-NT expression in our newly generated “humanized” *SCN5A* alt-pA knock-in mice; however, western blotting with the available antibodies failed to detect any convincing bands (data not shown); enrichment methods (e.g. immunoprecipitation) might be required. Overall, these data support that expression of transgene-derived NaV1.5-NT in cardiomyocytes can produce a detectable and stable protein both *in vitro* and *in vivo*.

**Figure 3.**
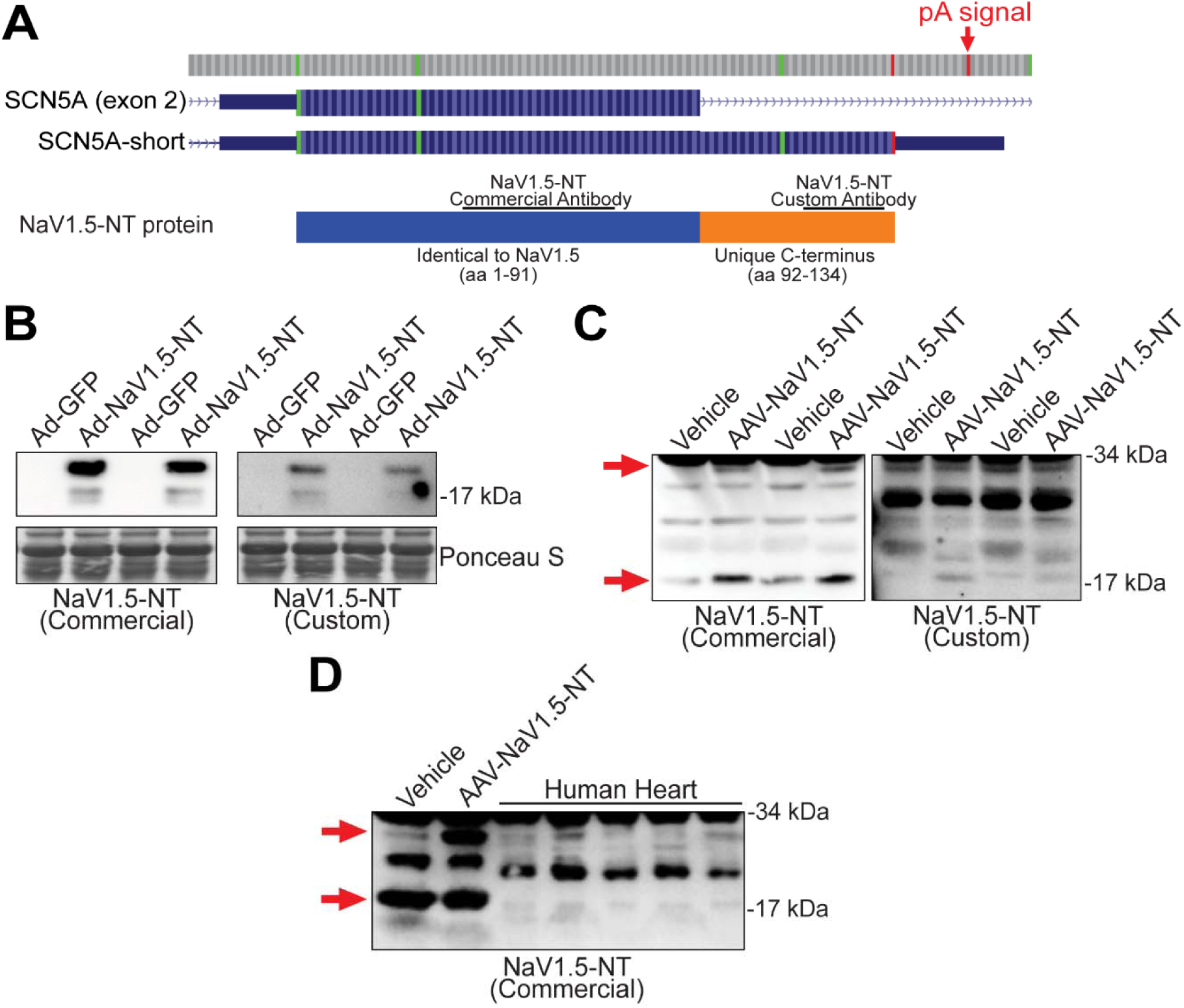
*SCN5A-short* generates a novel protein corresponding to the N-terminus of NaV1.5. A) Schematic of protein generated from *SCN5A-short*, which we call NaV1.5-NT, since it corresponds to the N-terminus of NaV1.5 (blue region; amino acids 1-91) and a unique C-terminal region corresponding to “intronic” sequence (orange region; amino acids 92-134). Two antibody epitopes are shown: a commercial NaV1.5 antibody (Aviva; recognizing amino acids 42-70) and a custom antibody that we generated (recognizing amino acids 109-123). **B)** Western blot of NRCMs transduced with Ad-GFP or Ad-NaV1.5-NT using the commercial or custom NaV1.5-NT antibody which show unique NaV1.5-NT-derived bands running near the expected size. **C)** Western blot of mouse hearts transduced with cardiomyocyte-specific AAV-GFP or AAV-NaV1.5-NT using the commercial or custom NaV1.5-NT antibody. The commercial antibody shows a band at 17 kDa and a band at 34 kDa and the custom antibody shows a band at 17 kDa. **D)** Western blot of human hearts alongside control AAV transduced mouse hearts using the commercial NaV1.5-NT antibody shows bands co-migrating with 34 kDa and 17 kDa NaV1.5-NT that are present in human samples.

To assess whether NaV1.5-NT is endogenously expressed in humans, we performed western blot on human heart tissue lysates, which revealed bands running at the same sizes (∼17 and 34 kDa) as a positive control (heart lysate from AAV-NaV1.5-NT injected mouse; **Fig. 3D**). Note: we routinely find that freeze-thaw of these mouse heart control lysates appears to alter the distribution of NaV1.5-NT (17 versus 34 kDa) and/or increases the prominence of the background band at 17 kDa, seen strongly now in both vehicle and AAV treated samples. To gain additional evidence of the presence of NaV1.5-NT in humans, we searched publicly available mass spectrometry data using PepQuery, a publicly available peptide identification algorithm^43^. This search yielded several peptides corresponding to the unique C-terminal region of NaV1.5-NT that were identified in human cells and tissues and scored as “confident”, providing further support of its natural existence in humans (**Table S1**).

### NaV1.5-NT localizes primarily to mitochondria

Next, we sought to gain insight into the molecular function of NaV1.5-NT, initially hypothesizing that it may influence NaV1.5 channel activity, based on a previously published report showing that transgene-mediated expression of an artificial NaV1.5 NT fragment (aa 1-132) was sufficient to enhance NaV1.5 current densities^44,45^. Along these lines, we tested if adenoviral expression of NaV1.5-NT in NRCMs altered NaV1.5 activity; however, we observed no difference in Na+ current densities under low extracellular Na+ concentration nor at physiologic Na+ concentration, the latter of which drives high Na+ influx; (**Supplementary** Fig. 2).

While examining the NaV1.5-NT amino acid sequence using various *in silico* prediction algorithms, we found that it contains a predicted (and possibly cleaved) mitochondrial targeting sequence (predicted by both MitoProtII^46^ and MitoFates^47^). To examine this further, we transfected NRCMs with NaV1.5-NT expression plasmid and performed immunofluorescence staining. NaV1.5-NT staining (not present in adjacent non-transfected cells) showed clear co-localization with mitochondrial staining (MitoTracker), which was abolished upon deletion of the predicted mitochondrial targeting sequence (ΔMTS; **Fig. 4A**). We next assessed the presence of NaV1.5-NT by western blotting of mitochondrial isolates collected from Ad-infected (NaV1.5-NT or GFP) NRCMs; this showed enrichment of two out of three NaV1.5-NT derived bands in mitochondrial lysates, evidenced by decreased signal in the mitochondria-depleted supernatant (Sup) samples and appearance in mitochondria pellets (**Fig. 4B**). Interestingly, the smallest band is not enriched in mitochondrial pellets and may derive from an alternative in-frame ATG codon located downstream of the mitochondrial targeting sequence. To further define the intra-mitochondrial localization of NaV1.5-NT, we performed classical proteinase K extraction of mitochondrial proteins. These data demonstrate that NaV1.5-NT resides within the mitochondrial matrix, consistent with the absence of a transmembrane domain (via Phobius^48^; **Fig. 4C**. In addition to NRCM studies we, blotted mitochondrial pellet lysates collected from AAV-treated mouse hearts, and the resulting data supports that NaV1.5-NT also localizes to mitochondria *in vivo* (**Supplementary** Fig. 3) Together, these data indicate that NaV1.5-NT is most likely a mitochondrial matrix protein.

**Figure 4.**
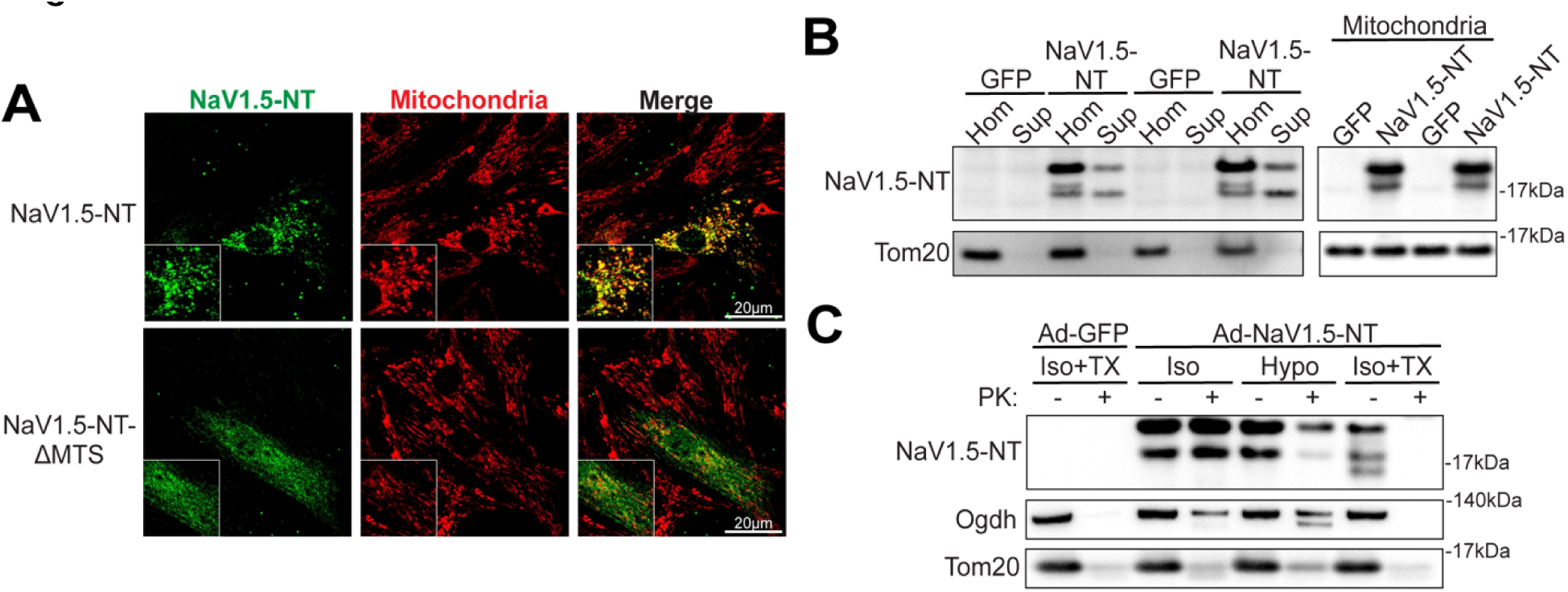
NaV1.5-NT localizes primarily to mitochondria. A) Confocal images of immunostained (commercial NaV1.5-NT antibody) NRCMs transfected with plasmids expressing NaV1.5-NT or NaV1.5-NT-ΔMTS (mitochondrial targeting sequence deleted). Co-localization with mitochondria (labeled with MitoTracker) was evaluated. Representative images are shown. Scale bar, 20µm. **B)** Mitochondria were isolated from NRCMs transduced with adenoviruses (Ad-GFP or Ad-NaV1.5-NT). Fractions corresponding to the total homogenate (Hom), mitochondria-depleted supernatant (Sup), or mitochondrial pellets were blotted for NaV1.5-NT (commercial antibody) and TOM20, a classic marker for mitochondria. **C)** NRCM mitochondria isolated from Ad-GFP or Ad-NaV1.5-NT transduced cells were treated with isotonic (Iso), hypotonic (Hypo), or isotonic plus triton (Iso+TX) in the presence or absence of proteinase K (PK) and subjected to western blot analysis. Proteins with known localization to the mitochondrial matrix (Ogdh) or outer membrane (Tom20) were blotted for as controls.

### NaV1.5-NT expression influences mitochondrial respiration

We next explored whether NaV1.5-NT functionally influences mitochondrial physiology. Given prior work suggesting that *SCN5A*/NaV1.5 may influence mitochondrial Ca2+ through crosstalk with the mitochondrial Na+/Ca2+ exchanger (Nclx)^12^, we assessed whether NaV1.5-NT expression might influence mitochondrial Ca2+ dynamics in NRCMs; however, we found no effects on mitochondrial Ca2+ levels nor influx/efflux rates (**Supplementary** Fig. 4). Next, we measured respiration in Ad-transduced NRCMs by Seahorse assay (mitochondrial stress test). Baseline oxygen consumption rates were clearly and consistently elevated in NaV1.5-NT transduced cells compared to multiple controls (untreated, Ad-GFP, or Ad-βGal; **Fig. 5A**). Notably, this increased basal respiration was inhibited by oligomycin, suggesting that NaV1.5-NT might promote proton leak through complex V. NaV1.5-NT also significantly increased maximal respiration, albeit to a lesser degree with some inconsistency across experiments (see extended data in **Supplementary** Fig. 5). In a complementary experiment (Seahorse ATP rate assay), similar findings were obtained, with NaV1.5-NT expression inducing a much higher rate of inferred mitochondrial (OXPHOS-derived) ATP production (**Fig. 5B, Supplementary** Fig. 6). We further examined this using a ratiometric ATP-sensitive GFP sensor^49^ and found that NaV1.5-NT expression increased ATP levels in NRCMs (**Fig. 5C**). Given that NaV1.5-NT increases mitochondrial respiration through OXPHOS, we examined if this is linked to elevated mitochondrial ROS and depleted NADH pools. We measured mitochondrial and cytosolic ROS using genetically encoded redox sensors^50^ and found that NaV1.5-NT expression significantly increased mitochondrial-derived ROS, with no effect on cytosolic ROS levels (**Fig. 5D, Supplementary** Fig. 7). In addition, we found that Ad-NaV1.5-NT decreased the amount of total NADH levels but had no effect on NAD+, resulting in overall increased NAD+/NADH ratio (**Fig. 5E**).

**Figure 5.**
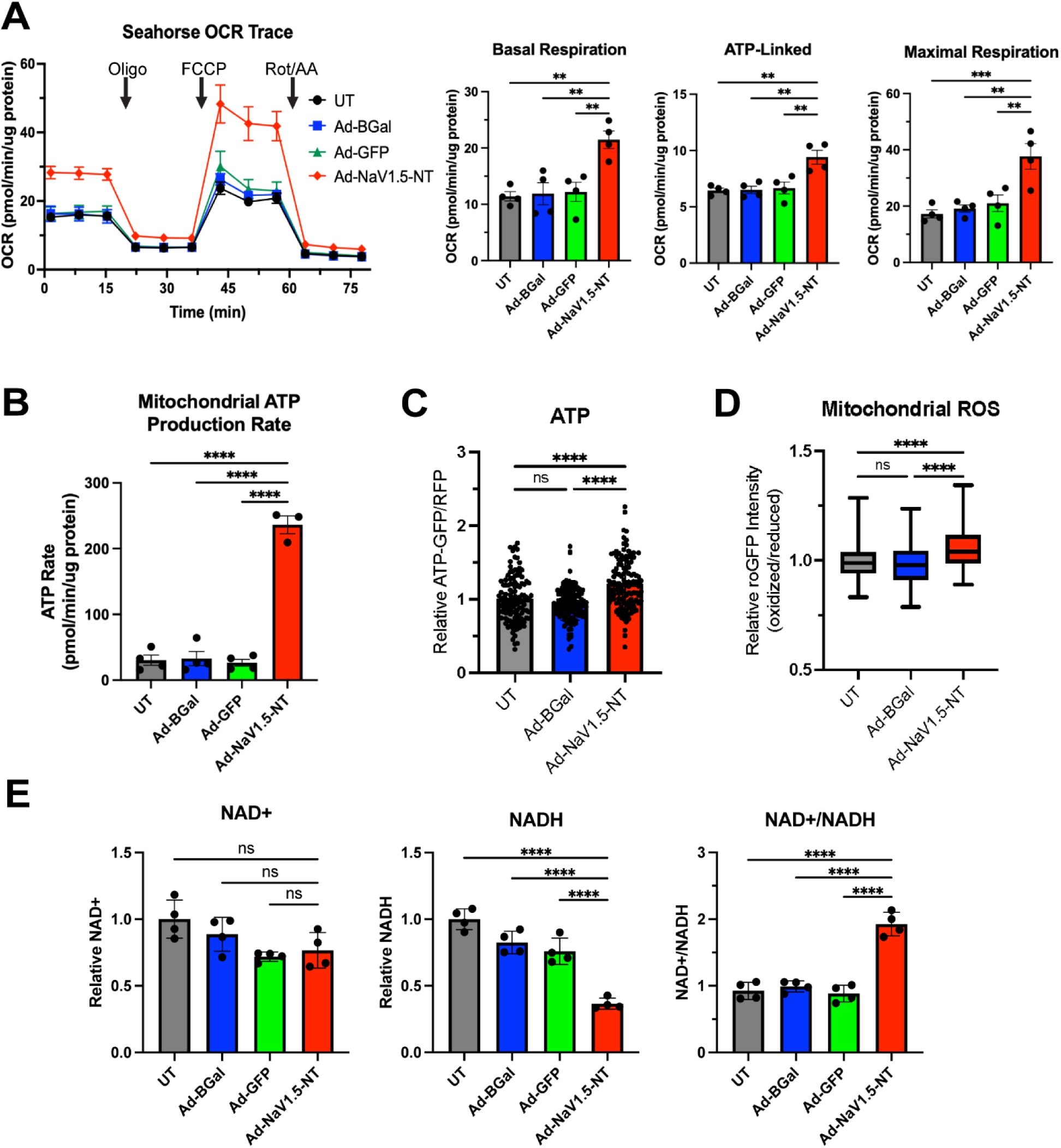
NaV1.5-NT expression influences mitochondrial respiration. A) Seahorse assay (mitochondrial stress test; 0.3um FCCP dose) of Ad-NaV1.5-NT transduced NRCMs (untreated, Ad-GFP, and Ad-βGal controls are also shown). Basal, ATP-linked, and maximal respiration are shown to the right and are plotted as mean ± SEM, p-values were determined by a one-way ANOVA with Dunnett’s post-hoc (**p<0.01, ***p<0.001). Refer to **Supplementary Fig. S5** for additional measures. **B)** Mitochondria-derived ATP production rates calculated from Seahorse ATP rate assay. Plotted as mean ± SEM, p-values were determined by a one-way ANOVA with Dunnett’s post-hoc (****p<0.0001). **C)** Relative ATP levels in Ad-transduced NRCMs, quantified by fluorescence intensity of a ratiometric ATP-sensitive GFP reporter. Plotted as mean ± SEM, p-values were determined by a one-way ANOVA with Tukey’s multiple comparison’s test (****p<0.0001). **D)** Mitochondrial ROS levels in Ad-transduced NRCMs, quantified by mitochondrial-localized ratiometric redox sensitive GFP sensor (Ad-roGFP). Minimum to maximum box and whisker plots are shown, p-values were determined by one-way ANOVA with Tukey’s multiple comparison’s test (****p<0.0001). **E)** NAD+ and NADH quantification in adenovirus transduced NRCMs. Plotted as mean ± SEM, p-values were determined by a one-way ANOVA with Tukey’s multiple comparison’s test (****p<0.0001).

### NaV1.5-NT increases complex I-containing supercomplexes and modulates complex V subunits

Given that NaV1.5-NT expression led to a striking decrease in NADH supply and increased baseline mitochondrial oxygen consumption rate, which was oligomycin-sensitive, we speculated that NaV1.5-NT may influence OXPHOS complexes, specifically complexes I and V respectively. To begin assessing this, we performed blue native-PAGE (BN-PAGE) and western blot on mitochondrial isolates collected from Ad-transduced (NaV1.5-NT or GFP control) NRCMs to measure levels of OXPHOS (super)complexes (antibody cocktail; **Fig. 6A**). We found no obvious differences in the amounts of primary OXPHOS complexes but noted the clear presence of several bands (migrating at ∼440 kDa) that are increased with NaV1.5-NT expression. Signal for intact complex I was weak and unclear given the high abundance of complex V, thus we ran additional gels and blotted specifically for complex I subunit Ndufb8 (**Fig. 6B**). This revealed that NaV1.5-NT expression increases the abundance of high molecular weight supercomplexes containing complex I, with no noticeable effect on the levels of primary complex I or total Ndufb8 expression (SDS-PAGE, **Supplementary** Fig. 8). Notably, this increase in supercomplexes was also reflected in in-gel complex I activity assays (**Fig. 6B**).

**Figure 6.**
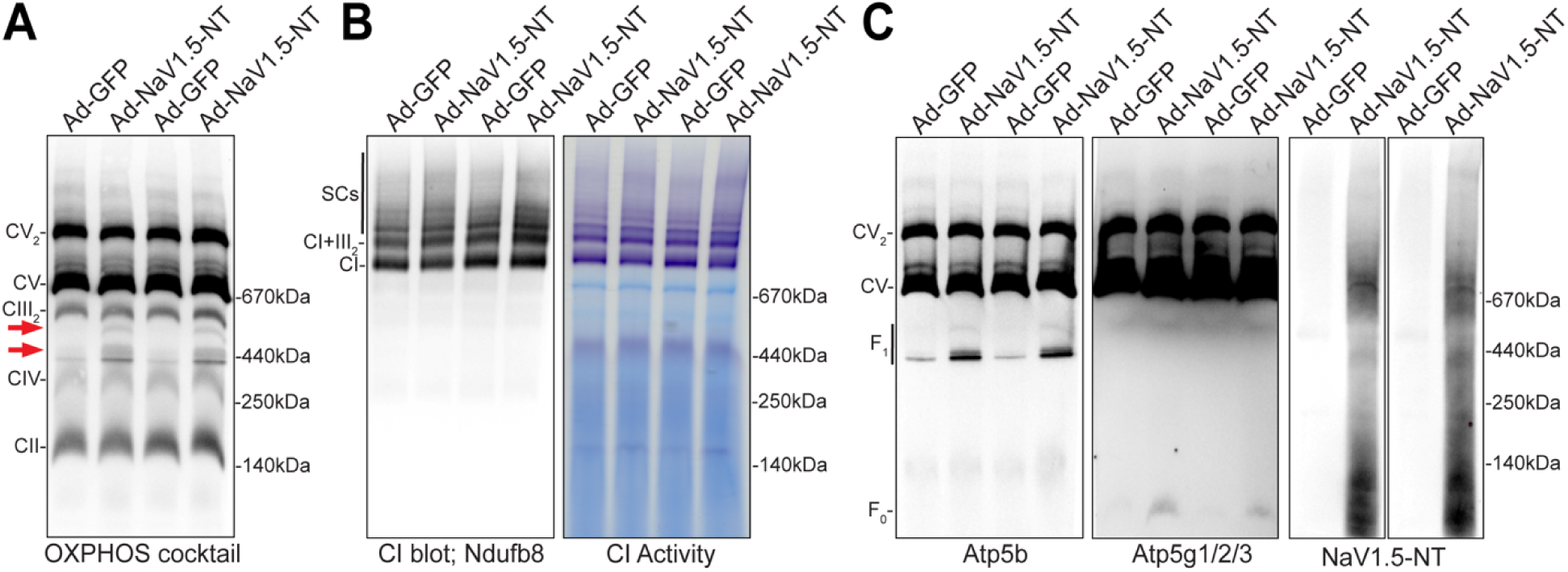
NaV1.5-NT influences complex I-containing supercomplexes and complex V subunits. A) Representative BN-PAGE blots of Ad-GFP and Ad-Nav1.5-NT treated NRCM mitochondrial isolates using an OXPHOS cocktail antibody. **B)** In Ad-GFP and Ad-NaV1.5-NT treated NRCM mitochondrial isolates, representative BN-PAGE blot using a complex I antibody (Ndufb8; left) or in-gel complex I activity (violet staining; right). **C)** Representative BN-PAGE blots of Ad-GFP and Ad-Nav1.5-NT treated NRCM mitochondrial isolates using an Atp5b (F_0_-subunit), Atp5g1/2/3 (F_1_-subunit), or NaV1.5-NT (commercial) antibody.

We next examined complex V more closely given that many papers have shown that bands in the 440-500 kDa range can correspond to the F_1_ portion and intermediates of the ATP synthase (complex V)^51,52^. Indeed, BN-PAGE blotting for Atp5b, an F_1_ subunit, clearly indicated that these bands of interest (i.e. those increased with NaV1.5-NT expression) correspond to F_1_ subcomplexes, suggesting that NaV1.5-NT influences complex V biogenesis and/or stability (**Fig. 6C**). In addition, subsequent BN-PAGE blotting for subunit c of the ATP synthase (Atp5g1/2/3), showed increased levels of free F_0_-ATPase (c-ring) after NaV1.5-NT expression, suggesting that NaV1.5-NT might promote F1-F0 disassembly (**Fig. 6C**). These data combined with our mitochondrial phenotyping data lead us to speculate that NaV1.5-NT plays a multifaceted role in influencing mitochondrial physiology. Our data support that 1) NaV1.5-NT increases baseline respiration likely by promoting complex V subcomplex conformations that enhance proton leak, and 2) NaV1.5-NT increases overall respiratory efficiency by enhancing formation and/or stability of complex I containing respiratory supercomplexes, though the specific molecular mechanisms underlying each of these remain unresolved.

## DISCUSSION

Through careful examination of genome annotation tracks and various RNA-sequencing datasets, we discovered that the *SCN5A* gene contains a well-conserved polyadenylation signal downstream of the second exon. This leads to the biosynthesis of a truncated transcript isoform, which is downregulated in failing human hearts, perhaps hinting at a relevant role in heart disease risk and/or pathogenesis. Notably, this short transcript isoform generates a protein homologous to the N-terminus of NaV1.5, with a unique C-terminus, which we called NaV1.5-NT. This protein appears to be stable in isolated cardiomyocytes and in mouse hearts, the latter of which shows two bands running at 17kDa and 34kDa, perhaps an SDS-resistant dimer. The presence of NaV1.5-NT in humans was supported by western blot on human cardiac tissue samples, which was bolstered by the detection of several peptides identified in publicly available mass spectrometry from human cells and tissues. While this evidence is highly supportive of the natural existence of NaV1.5-NT in human tissue, future studies will need to implement high-resolution mass spectrometry of human heart tissues with proper control peptides.

NaV1.5-NT localizes to the mitochondrial matrix where it influences mitochondrial respiration. Baseline mitochondrial oxygen consumption was strikingly elevated, with increases in ATP-linked and maximal respiration upon NaV1.5-NT expression in NRCMs. BN-PAGE analyses of OXPHOS complexes showed increased levels of disassembled complex V. This likely leads to elevated proton leak via the low-conductance pore forming F_0_ subunit, resulting in enhanced oxygen consumption^53^ that is still inhibitable with oligomycin, which can bind to the F_0_ pore to inhibit proton transport^54^. Ultimately, future experiments aimed at measuring mitochondrial membrane potential and/or electrophysiological recordings of isolated mitoplasts will be needed to further define whether NaV1.5-NT expression is leading to proton leak via disassembled complex V.

Concurrently, we found that NaV1.5-NT expression increased levels of complex I-containing supercomplexes. This may be a secondary effect of proton leak, whereby complex I compensates to re-establish proper membrane potential. Another explanation is a multifaceted role for NaV1.5-NT in regulating complex V and complex I supercomplex (dis)assembly and/or stability. Indeed, our BN-PAGE analyses show that NaV1.5-NT appears to co-migrate near intact complex I and complex V subunits (**Fig. 6C**), however, co-immunoprecipitation studies will be needed to identify specific NaV1.5-NT protein interactors.

In summary, we serendipitously discovered a novel APA signal in *SCN5A*, which regulates full-length *SCN5A* mRNA expression and leads to the biosynthesis of a truncated transcript isoform and an N-terminal fragment of NaV1.5 that localizes to mitochondria where it influences respiration via complex I and/or complex V. Future studies in cultured cardiomyocytes (human iPS-derived) and in our *SCN5A* alt-pA-KI mice will need to examine how *SCN5A* APA is regulated or influenced by cardiac stress conditions. In addition, follow-up experiments will need to corroborate our findings *in vivo*, by expressing NaV1.5-NT in mouse hearts and performing cardiac myofiber respiration studies, mitochondrial phenotyping, and possibly metabolomics, which may give additional insights into underlying mechanisms. Additional studies will also be needed to determine if NaV1.5-NT expression influences disease outcomes in animal models of cardiac stress to begin exploring its potential translational relevance.

## ACKNOWLEDGEMENTS

This work was supported by the University of Iowa Carver College of Medicine (Distinguished Scholars Program to R.L.B.), NIH NHLBI (HL144717, and HL150557 to R.L.B.), NIH NIGMS (predoctoral fellowship T32 GM067795 to N.H.W. and undergraduate fellowship T34GM141143 to J.M.H.), American Heart Association (20IPA35360150 to R.L.B. and 23PRE1011277 to N.H.W.), University of Iowa College of Liberal Arts and Sciences (fellowship to J.M.H.). We acknowledge the University of Iowa core facilities that made significant contributions to this work, including the Genome Editing Facility, Cardiovascular Phenotyping Core, Viral Vector Core, and Free Radical and Radiation Biology. We also acknowledge Jasmyn Hoeger, Connor Linzer, and Jason Schwarzhoff for their assistance with plasmid cloning and other general project support, as well as the laboratories of Barry L. London, Ethan J. Anderson, and Isabella M. Grumbach for their technical assistance, scientific discussions, and resource sharing.

## AUTHOR CONTRIBUTIONS

R.L.B. conceived the project, supervised research and analyzed and interpreted data. N.H.W., J.M.M., C.S.S., J.M.H. designed and executed experiments, curated and analyzed data, and participated in data interpretation. J.Y.Y. and E.B. executed experiments and analyzed data. J.W.E. and B.L.L. guided experimental design, supported execution, and participated in data interpretation. N.H.W. and R.L.B. wrote the manuscript.

## SUPPLEMENTAL INFORMATION

Document S1. Figures S1-8, Table S1.

## MATERIALS AND METHODS

### Bioinformatics

*RNA-sequencing data visualization (related to Fig. 1A):* Human arial and left ventricular RNA-seq tracks were visualized on the UCSC Genome Browser^39^ (GTEx RNA-seq track^55^). For AC16, RH30, and iPS-cardiomyocytes, raw RNA-seq fastq files were downloaded from the SRA archive or European Nucleotide Archive (SRP131497, SRP323523, and PRJEB25366, respectively). Reads were aligned to a human reference genome (hg38) using STAR aligner (version 2.7.6), using the recommended settings, to generate bam files and were viewed on the Interactive Genomics Viewer (version 2.13.2).

*3’-end sequencing polyA usage analysis:* Processed human heart 3’-end polyA sequencing data was visualized on the UCSC Genome Browser using a link provided by the study authors. Peak heights corresponding to the upstream (*SCN5A-short*) polyA site and full-length *SCN5A* were manually tabulated and the ratio of upstream to full-length peak heights were calculated.

*SCN5A transcript quantification in diseased human hearts:* Fastq files were downloaded from SRA archive (SRP151309 and SRP237337). Fasta sequences for human *SCN5A-short*, full-length *SCN5A,* or *GAPDH* 3’-UTRs used to build a custom kallisto index. Raw RNA-seq fastq files were aligned to the index using kallisto with 100 bootstratps in single-end or paired-end (depending on the dataset) mode assuming the fragment length distribution is truncated gaussian with mean = 200 and sd = 30. Transcript per million (TPM) were calculated for each sample, then normalized to GAPDH TPM, and then the ratio of short / long was calculated per sample.

*PepQuery mass-spectrometry search*: PepQuery version 2.0.2 wash used for this analysis using the recommended settings to search all mass spec data sets in the PepQuery Database, using the Gencode human proteome as the reference data set. “protein” mode was used to search for spectra matching the unique C-terminal region (derived from “intronic” sequence) of human NaV1.5-NT: VTTTHLQPCLPFCATPLSMEQRGKRAWPPYGALFRVAHEALGK.

### Plasmids and Viral Vectors

The NaV1.5-NT open reading frame was PCR amplified from human heart cDNA using primers with added restriction enzyme sites and clones into viral CMV-based expression plasmids (University of Iowa Viral Vector Core). Primers sequences are provided in **Table S2**. AAV shuttle plasmids were generated by cloning NaV1.5-NT or GFP into a custom cardiac-specific vector (hsTNNT2-MCS-miR122b-WPRE-bGHpA). To achieve cardiomyocyte specificity, we used the human TNNT2 promoter and included miR122b binding sites to de-target the liver. AAV vectors were prepared by the University of Iowa Viral Vector Core and were packaged into AAV2/9 serotype capsids by standard triple transfection method. AAV purification was done with an iodixanol gradient followed by ion exchange into biocompatible F68/PBS storage buffer and titers (vg/mL) were determined by ddPCR. CMV-based adenoviral vectors were generated the University of Iowa Viral Vector Core using the RapAdI system^56^: Ad-BGal (Iowa-3554, Ad5CMVcytoLacZ), Ad-GFP (Iowa-4, Ad5CMVeGFP), Ad-NaV1.5-NT (Iowa-custom, Ad5CMVNaV1.5-NT). The cytosolic ATP, cytosolic redox, and mitochondrial redox sensors were obtained from Addgene (plasmid # 102551, 64991, and 64992, respectively) and cloned into an adenoviral shuttle plasmid before virus generation. For adenoviruses, approximate relative titers were determined by hexon immunostaining in HEK293T cells.

### Mouse Studies

All animal studies were approved by the Institutional Animal Care and Use Committees (IACUC) at the University of Iowa, and all experiments conform to the appropriate regulatory standards. Rodents were housed in a controlled temperature environment on a 12 hr light/dark cycle, with food and water provided ad libitum. Animal age and sex are noted within the manuscript where appropriate.

*SCN5A-altpA-KI mouse line:* CRISPR/Cas9-based gene editing was performed via the University of Iowa Genome Editing Facility.

*AAV Injection:* 5.2e10vg/g of purified AAV particles (AAV2/9-hsTNNT2-NaV1.5-NT-miR122b-WPRE-bGHpA or AAV2/9-hsTNNT2-eGFP-miR122b-WPRE-bGHpA) or vehicle buffer (F68/PBS) were injected intraperitoneally in 10-day old c57BL/6NJ (JAX-005304) mice. 6 weeks post-injection, mice were anesthetized under isoflurane and the heart was excised, washed in ice-cold saline, and snap frozen in liquid nitrogen.

### Cell Culture

*Neonatal Rat Cardiomyocytes:* NRCMs were isolated as previously described^57^ from 2-3 day-old rat pups (Sprague-Dawley, Charles River Stock 001). Cells were cultured in growth media (DMEM/F12 supplemented with 5% horse serum, 10mM HEPES, 1% ITS, 15ug/mL gentamycin, and 100uM BRDU), with daily media changes, and incubated at 37°C with 5% CO_2_.

*Plasmid Transfection:* The day after plating, NRCMs were transfected using 1ul Lipofectamine 2000 and 250ng plasmid DNA. Transfection complexes were formed in 50ul Opti-MEM, then diluted 4-fold and overlayed onto cells. After 4 hours incubation, cells were returned to growth media and incubated for 48-72 hours before assay.

*Adenoviral Transduction:* Crude adenoviral supernatant was diluted into growth media and added to cells the day after plating at ∼50 infectious units per cell. Media was changed 24 hours post transduction. Cells were assayed at 48-72 hours post-transduction, with daily media changes.

### RT-qPCR

Snap-frozen mouse heart tissue was crushed under liquid nitrogen. 50mg of tissue was added to ice-cold QIAzol lysis reagent, homogenized by beat beating at 30Hz for 2 minutes, and clarified by centrifugation at 10,000xg for 10minutes. Total RNA was isolated from the resulting supernatant using the Qiagen miRNeasy Mini Kit and gDNA was removed by the on-column DNAse treatment, according to the manufacturer’s directions. 1ug RNA was converted to cDNA using the Applied Biosystems High Capacity cDNA Reverse Transcription Kit, using oligo-dT primers. RT-qPCR was performed using custom-designed SYBR Green assays, done in triplicate and detected on a Viia 7 qPCR machine (Applied Biosystems). Standard curve and melt curves were evaluated for quality control and relative mRNA expression was determine by the ddCt method. Primers are listed in **Table S2**.

### SDS-PAGE and Western Blotting

Heart tissue protein lysates were prepared, as previously described^58^, by addition of lysis buffer to snap-frozen and crushed tissue, and disrupted by beat beating. NRCM cell lysates were prepared by addition of RIPA buffer to the cells, incubated on ice for 10 minutes, and transferred to tubes. Protein extracts were clarified by centrifugation at 10,000xg for 10mins and quantified by BCA assay. Protein concentration was equilibrated between samples and sample buffer was added (NuPAGE reagents). Proteins were resolved by standard SDS-PAGE on 4-12% Bis-Tris gels using MES buffer (ThermoFisher) before being transferred to 0.2micron nitrocellulose membranes. The membrane was blocked in 5% milk in TBST, incubated with the indicated primary antibody overnight at 4°C, washed, incubated with HRP-conjugated secondary antibody for 1hour at room temperature, and washed. Proteins were visualized by chemiluminescence using West Femto Maximum Sensitivity Substrate (ThermoFisher) and imaged using an iBright 1500.

### Immunofluorescence Staining and Confocal Microscopy

48 hours post-transfection, mitochondria were labeled in live cells by incubation with 100nM MitoTracker Deep-Red FM (ThermoFisher) in media for 30min. Cells were wash 3 times, 10mins each with media and fixed with 4% paraformaldehyde for 10mins at 37°C. For immunostaining, cells were washed with PBS and blocked/permeablized in PBS containing 5% goat serum and 0.2% Triton-X100 rocking for 1hour at room temperature. Primary antibody (NaV1.5-NT commercial antibody) diluted in 10% blocking buffer in PBS and incubated rocking overnight at 4°C. Cells were washed in PBS and incubated in Alexa568 conjugated goat-anti-rabbit secondary antibody for 2 hours at room temperature and wash with PBS. Images were captured using a confocal laser scanning microscope (Zeiss LSM 510 meta) with the Zeiss ZEN software.

### Mitochondria Isolation

*From heart tissue:* All steps were performed at 4°C using ice-cold reagents. Freshly excised mouse heart was split in half and washed in ice-cold PBS and minced in a small volume of isolation buffer (250mM sucrose, 5mM Tri-HCl, 2mM EGTA, 5mM MgCl_2_, 1mM ATP, and 0.5% fatty acid-free BSA, pH 7.4). Minced tissue was washed with fresh isolation buffer once and homogenized manually using an 8mL Potter Elvehjem tissue grinder with PTFE pestle (Kimble 886000-022), in 4-6mL isolation buffer with 0.2% BSA for 6-8 passes. Cell debris was pelleted at 800xg for 10mins and the clarified homogenate centrifuged at 8000xg for 10min to pellet the mitochondria. Mitochondria pellets were snap frozen in liquid nitrogen and stored at -80°C until processing for downstream analyses.

*From NRCMs:* Method adapted from Frezza et al.^59^. Mitochondria were isolated from one 10cm dish per sample and all steps were performed at 4°C using ice-cold reagents. Cells were washed 2x with PBS and scraped in 2mL of isolation buffer ((200mM sucrose, 10mM Tris, 1mM EGTA, pH 7.4) and pellets at 800xg for 1min. Cells were resuspended in 1ml isolation buffer and manually homogenized in a 2ml dounce tissue grinder (Kimble) with 20 passes using “pestle B”. Cell debris and nuclei were removed by centrifugation at 800xg for 5mins. The resulting supernatant (Hom) was centrifuged at 8,000xg for 10mins to pellet mitochondria, the resulting mitochondria-depleted supernatant (Sup) was saved for some analyses. To test NaV1.5-NT enrichment in mitochondria, all fractions samples were protein-normalized to the Hom fraction. For all other experiments, samples were protein normalized to mitochondria.

*Proteinase K assessment of mitochondria proteins:* 48-hours post transduction, NRCM mitochondria were isolated from 2 10cm dishes for each sample and equally divided into 6 tubes. Mitochondria pellets were resuspended in either isotonic (isolation buffer), hypotonic (5 mM HEPES adjusted to pH 7.4 with KOH), or isotonic with 1% Triton X-100 buffer. 20ug/ml proteinase K was added to the indicated samples and all were incubated at room temperature for 20min. Proteins were TCA precipitated to deactivate the proteinase K by addition of trichloroacetic acid to a final concentration of 10% and incubated on ice for 1 hour. Protein precipitates were pelleted and washed twice with acetone before resuspension on sample buffer.

### Seahorse Assay

48 hours post-transduction, NRCMs were changed to oxygen consumption rate (OCR) measurements at 37°C in an XF96 extracellular flux analyzer (Seahorse Bioscience). Cells (3x104) were plated in XF media pH 7.4 supplemented with 5mM HEPES, 10mM glucose, 2mM L-glutamine, and 1mM sodium pyruvate. For the mitochondrial stress test, cells were sequentially exposed to oligomycin (2μM), FCCP (0.5μM), and rotenone plus antimycin A (1μM each). For the ATP production rate assay, cells were sequentially exposed to oligomycin (2μM) and rotenone plus antimycin A (1μM each). ATP rates were calculated using the Agilent Seahorse ATP rate calculator using experimentally determined buffer factor calculated during the experiment. All values were normalized to total protein determined for each well by BCA assay immediately after assay.

### NAD+/NADH Quantification

NAD+/NADH was measured as previously described^19^. 48-hours post-transduction, NRCMs plated in 24-well plates were washed with ice-cold PBS and subject to NAD+/NADH-Glo Assay (Promega, G9071), according to the manufacturers directions. Briefly, cells were lysed and split into two tubes – one for NAD+ and one for NADH measurement. NAD+ samples were acidified and heated to destroy NADH, and NADH samples were heated in the basic lysis buffer to degrade NAD+. Samples were neutralized and subject to NAD+/NADH assay to quantify each separately for each sample. Luminescence was recorded on a Synergy H1 microplate reader (BioTek) and values during the mid-linear phase of the reaction were selected for analysis and normalized to total protein amount.

### Blue Native (BN)-PAGE Blotting and cI Activity

Methods were performed as previously described^60^. Briefly, NRCM mitochondrial pellets were digitonin solubilized and 10ug protein was run on 3-12% Bis-Tris NativePAGE gels (ThermoFisher) using “dark blue” running buffer and changed to “light blue” running buffer about 1/3 of the way through. Protein complexes were then transferred to PVDF membranes using NuPAGE transfer buffer. The membrane was fixed in 10% acetic acid for 10mins, allowed to dry, washed in methanol until to remove the blue dye, and equilibrated with water. The membrane was then treated according to standard western blot procedures. For in-gel complex I activity, 30ug protein was run on Native-PAGE gels using “light blue” followed by “no blue” running buffer. The gel was immediately incubated in freshly prepared CI substrate solution (2 mM Tris-HCl pH 7.4, 0.1 mg/ml NADH, 2.5 mg/ml Nitrotetrazolium Blue chloride (NTB)) for 20-30 min on a rocker at room temperature. Appearance of violet bands indicates CI activity. Reaction was stopped by incubation in 10% acetic acid for 10min and washed in water.

---

**Figure S1:**
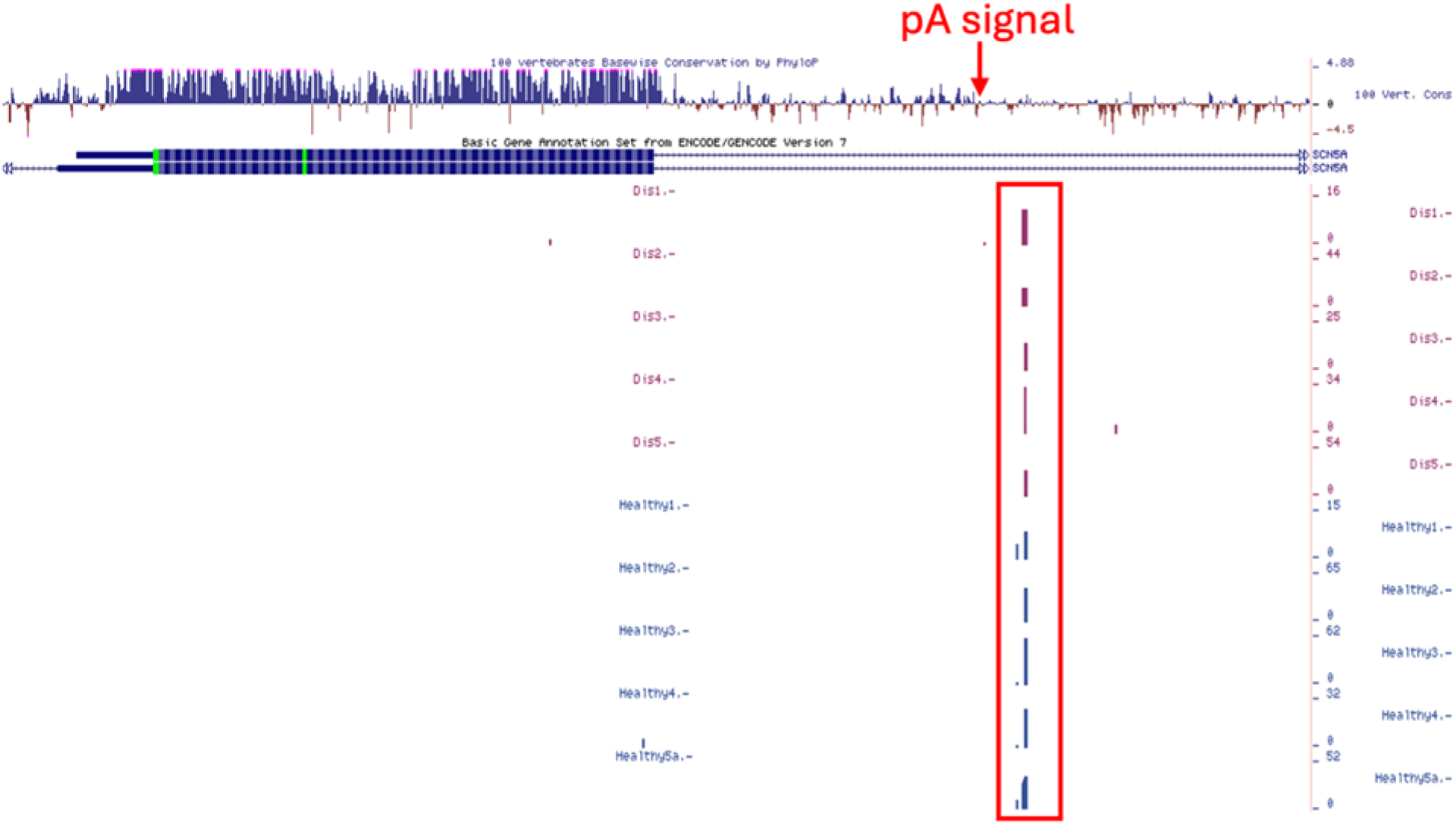
Human heart alternative polyadenylation maps show usage of a polyA signal downstream of *SCN5A* exon 2. Screenshot of the *SCN5A* exon 2 locus with tracks corresponding to alternative polyA maps from 10 human heart samples, all showing clear signal corresponding to a polyA signal sequence downstream of *SCN5A* exon 2. Red box indicates 3’-end sequencing signal, red arrow indicates the polyA signal sequence [AATAAA]).

**Figure S2:**
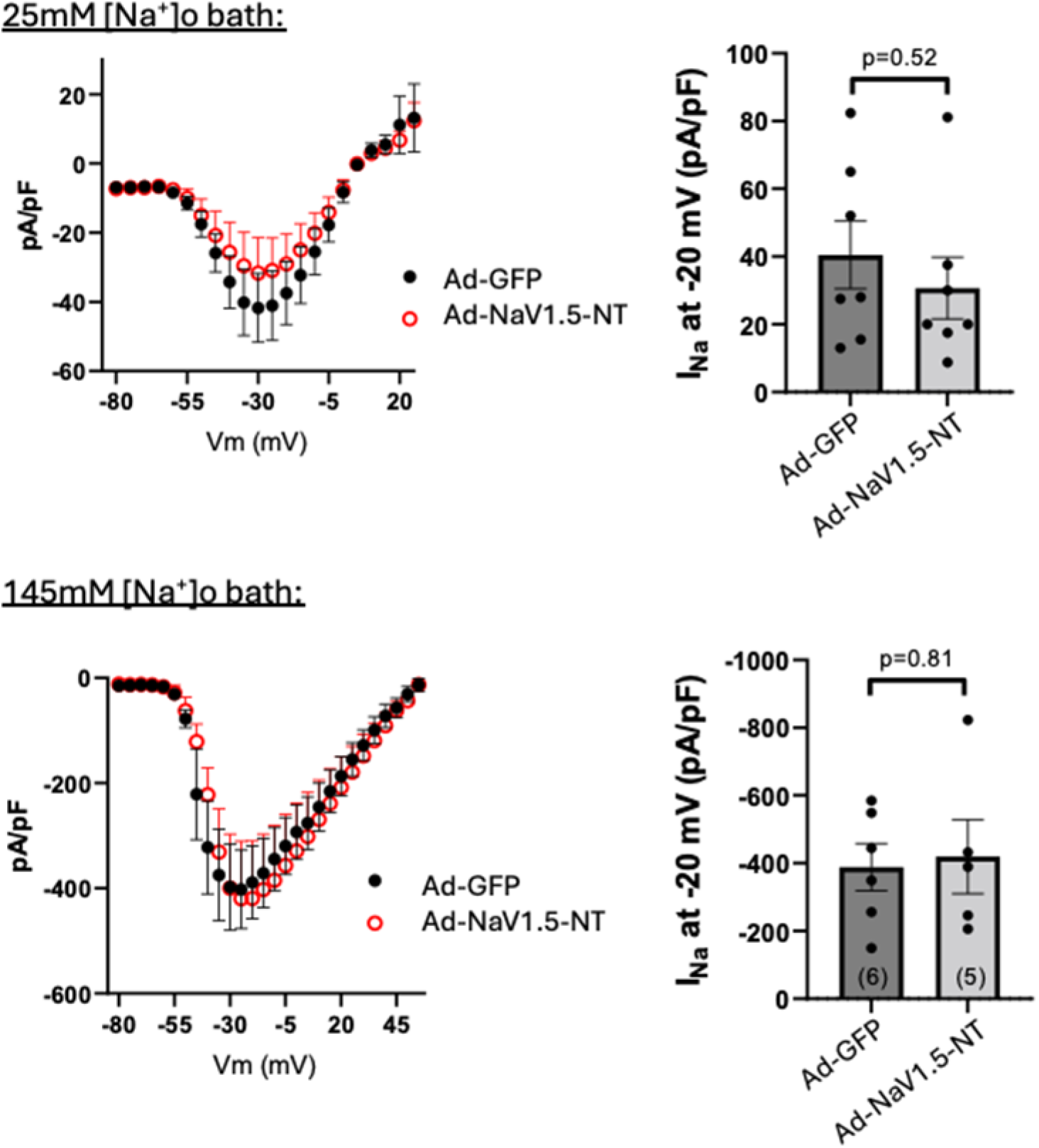
NaV1.5-NT does not alter sodium current in NRCMs. Patch-clamp electrophysiological recordings of Na+ currents in NRCMs treated with Ad-NaV1.5-NT or Ad-GFP (control) under low (25mM; top) or physiologic (145mM; bottom) extracellular Na+ concentrations. Peak Na+ currents are plotted to the right for each condition, plotted at mean ± SEM, and p-values were determined by an unpaired t-test.

**Figure S3:**
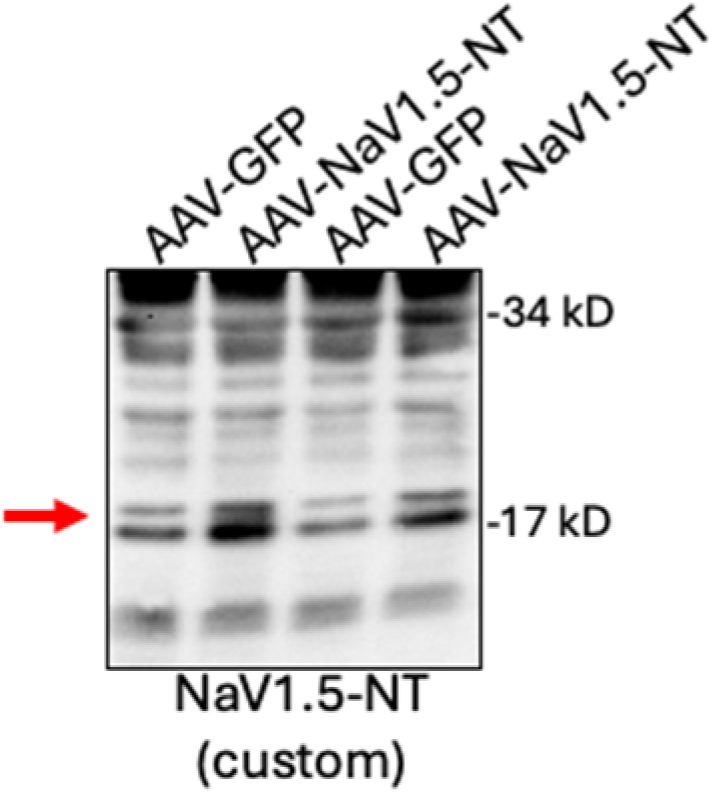
NaV1.5-NT is enriched in mouse heart mitochondria. Mouse hearts were transduced with AAV-GFP (control) or AAV-NaV1.5-NT, and mitochondria were isolated and subjected to western blot analysis for NaV1.5-NT (custom antibody). Red arrow indicates NaV1.5-NT band that is detected in these mitochondrial isolates.

**Figure S4:**
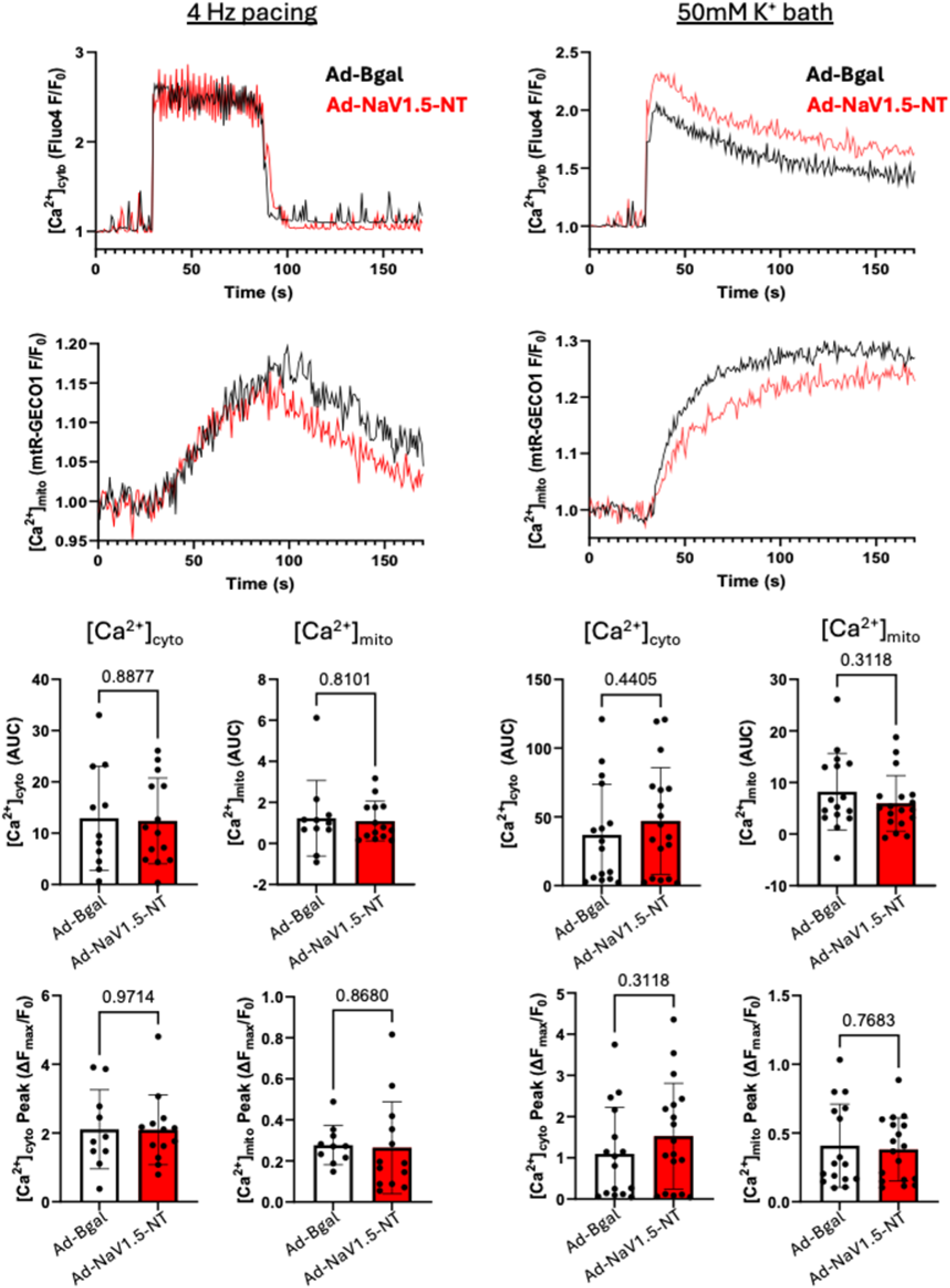
NaV1.5-NT does not alter cytosolic or mitochondrial calcium levels. Representative mitochondrial or cytosolic calcium traces recorded in Ad-treated NRCMs (BGal versus NaV1.5-NT) under 4Hz pacing (left) or 50mM K^+^ bath (right). Area under the curve (AUC) and peak calcium levels are quantified below for cytosolic and mitochondrial calcium. Data are plotted at mean ± SD, and p-values were determined by an unpaired t-test.

**Figure S5:**
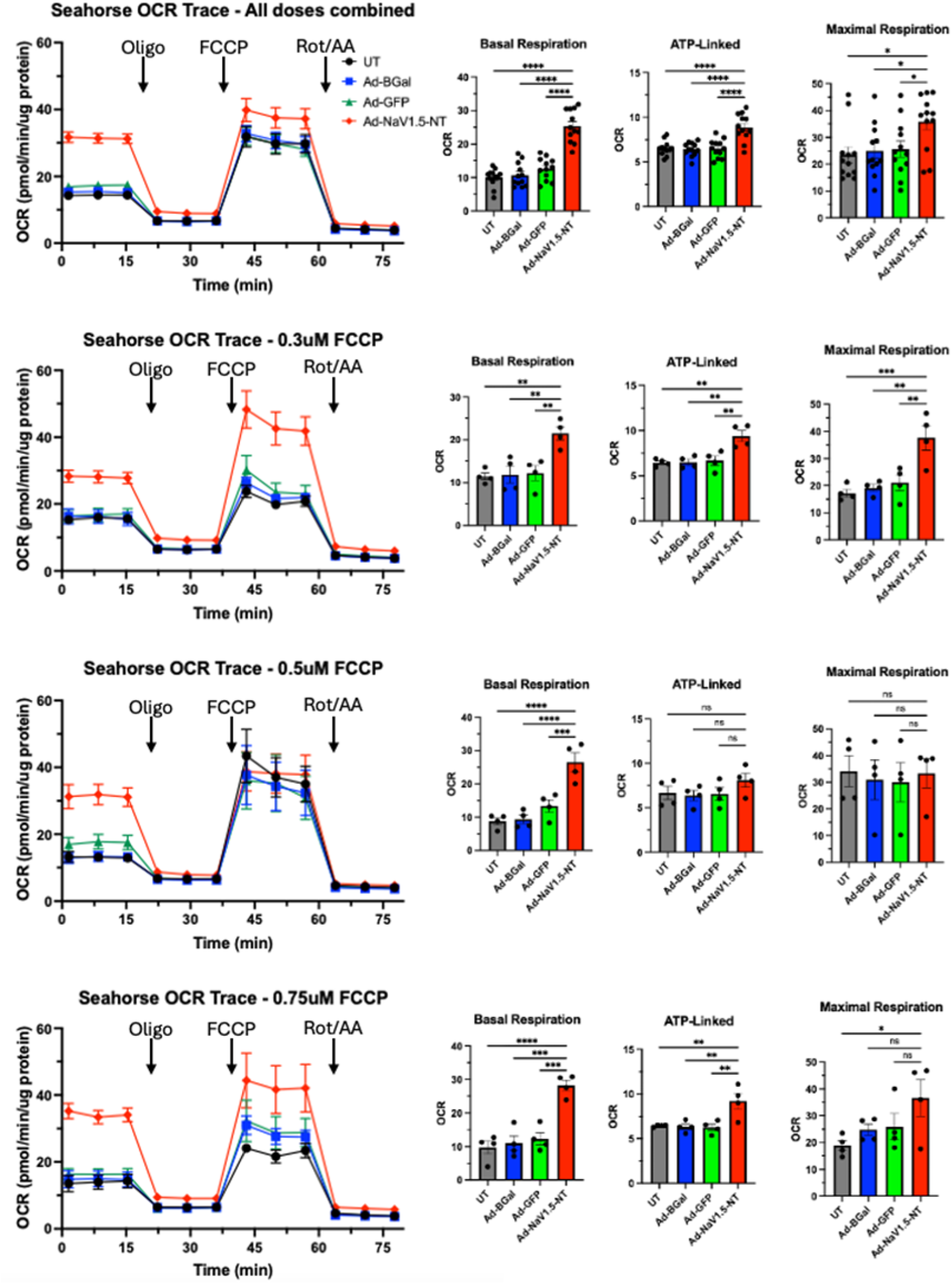
NaV1.5-NT alters mitochondrial respiration. Seahorse mitochondrial stress test in Ad-NaV1.5-NT or control (untreated, Ad-BGal, or Ad-GFP) treated NRCMs. Cells were treated with various concentrations of FCCP during the second injection. The compiled data are shown along with the data separated by FCCP concentration. Seahorse traces are shown to the left and basal, ATP-linked, and maximal respiration plots are shown to the right. Data are plotted as mean ± SEM, p-values were determined by a one-way ANOVA with Dunnett’s post-hoc (*p<0.05, **p<0.01, ***p<0.001, ****p<0.0001).

**Figure S6:**
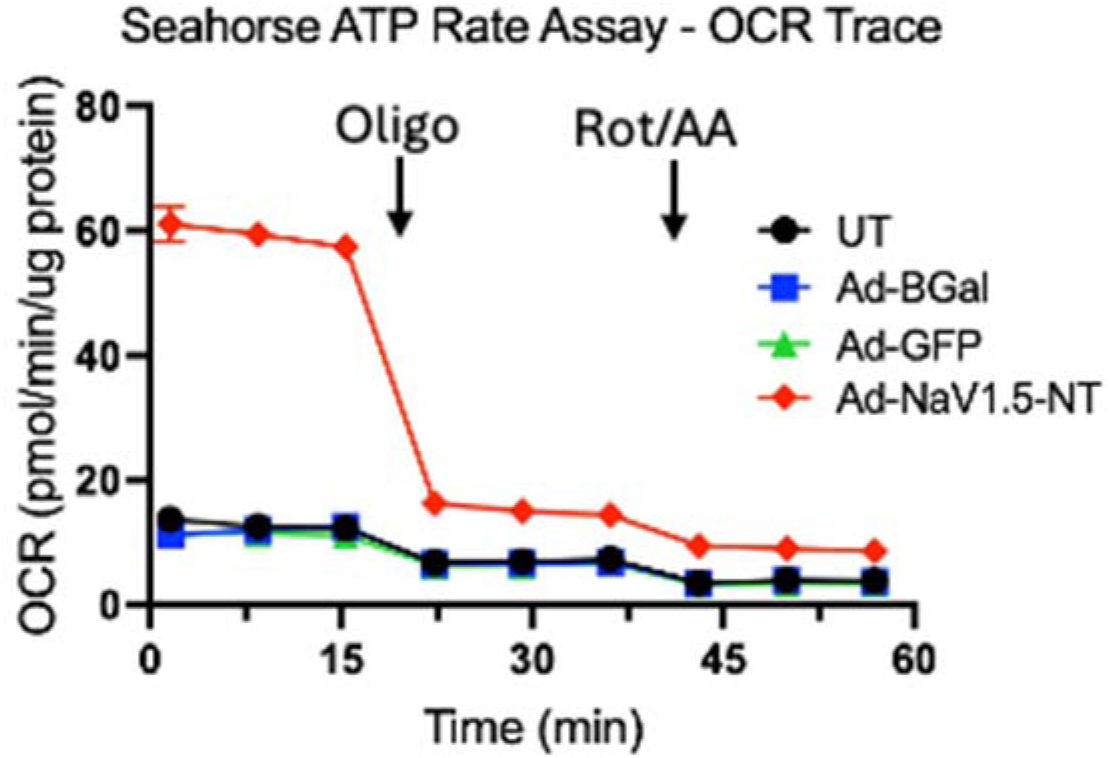
Seahorse ATP rate assay. Seahorse trace from the ATP rate assay, used to calculate mitochondrial ATP production rate in Ad-treated NRCMs (NaV1.5-NT versus various controls).

**Figure S7:**
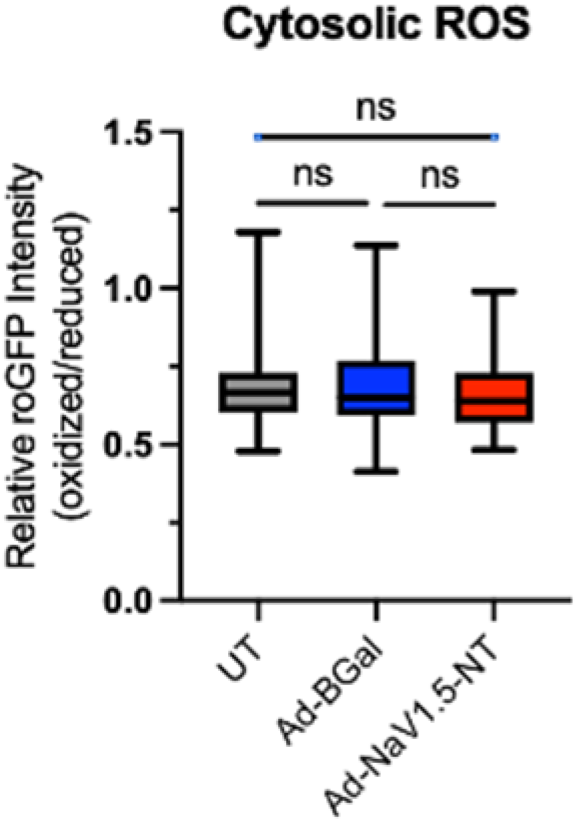
Cytosolic ROS levels. Cytosolic ROS comparison in Ad-treated NRCMs (NaV1.5-NT versus untreated or BGal controls) using a non-targeted ratiometric redox sensor. Minimum to maximum box and whisker plots are shown, p-values were determined by one-way ANOVA with Dunnett’s post-hoc.

**Figure S8:**
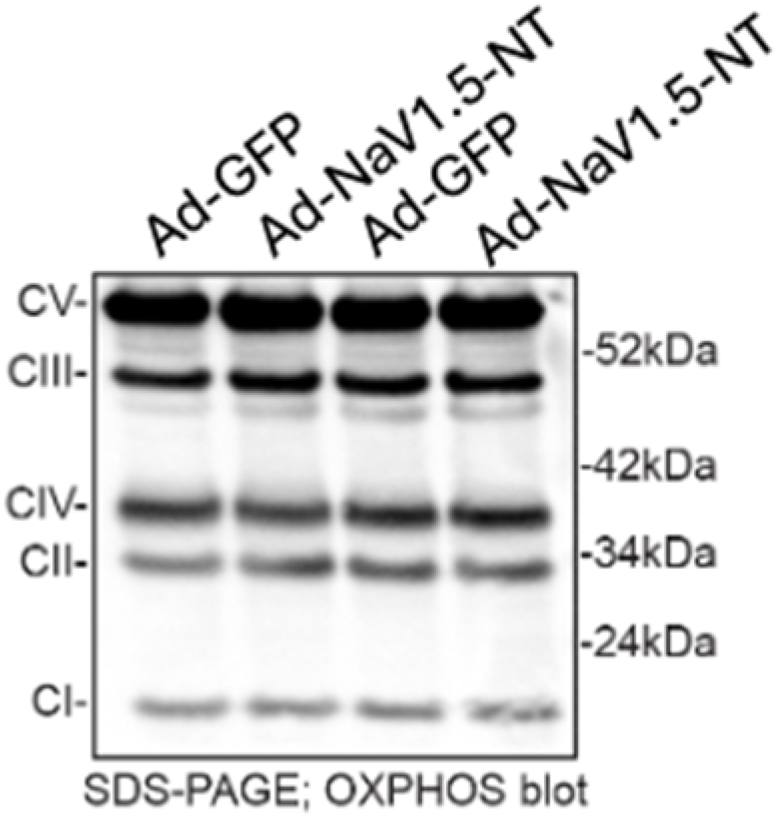
NaV1.5-NT does not alter representative OXPHOS complex subunit expression level. Western blot analysis of mitochondria isolated from Ad-treated (NaV1.5-NT versus GFP) NRCMs using an OXPHOS cocktail of antibodies.

**Table S1:**
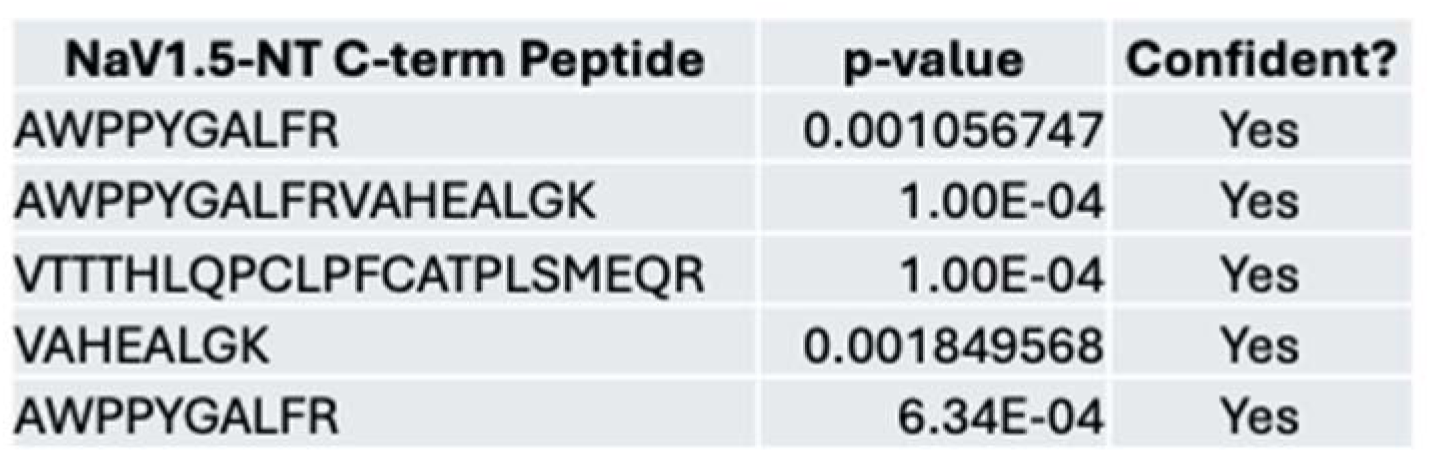
PepQuery search of human mass spectrometry data. Search of PepQuery datasets for peptides matching the unique C-terminal region of NaV1.5-NT. Only peptides scored as “confident” (i.e. significant) are shown.

